# Double-check the zebrafish *18s rRNA* qPCR primers: they may be wrong

**DOI:** 10.1101/2021.10.23.464780

**Authors:** Jianing Wang, Zhipeng Yang, An Xiao

## Abstract

A widely used qPCR primer for zebrafish gene *rna18s* (*18s rRNA*, or *18s*), with the sequence of 5’-TCGC<u>**ta**</u>GT<u>**t**</u>GGCATCGTTTA<u>**t**</u>G-3’, is found to be incorrect. Initially designed for rainbow trout (*Oncorhynchus mykiss*) *rna18s*, the primer has four different nucleotides from the zebrafish sequence 5’-TCGC<u>**GG**</u>GT<u>**C**</u>GGCATCGTTTA<u>**C**</u>G-3’ (indicated in bold/underlined, lowercase letters for rainbow trout and uppercase letters for zebrafish). Since its first use in zebrafish in 2006, this mismatched primer has been clearly stated to be used in at least 50 publications and may have affected hundreds or more in publications citing them. For a sensitive, quantitative method as qPCR, this error must be corrected as soon as possible in the zebrafish community by using *rna18s* primer sets with accurate sequences, such as those summarized and newly designed in this article.

## Introduction

Quantitative comparison of gene expression levels is critical for biological research. It is one of the basic approaches for exploring and analyzing physiology, pathology, and drug mechanisms in experimental organisms. The gene expression profiling studying approaches include Northern blotting, cDNA microarrays, RNA-seq, quantitative reverse transcription-polymerase chain reaction (RT-qPCR), serial analysis of gene expression (SAGE), etc. (Zhang et al., 2019). Among them, RT-qPCR is the most widely used assay to evaluate mRNA expression levels in different samples or experimental groups (Bustin, 2002). Due to the variation in the starting material, experimental steps of RNA extraction, and cDNA preparation, it is difficult to determine the absolute quantities of mRNA in the samples. Instead, relative quantification methods are employed to compare the expression of target genes by comparing their C_q_ (quantification cycle, also known as C_t_, threshold cycle) values in qPCR experiments to those of an internal control reference gene from equal amounts of the same PCR templates, assuming that the reference genes have similar expression levels among all samples and groups (Rebouças et al., 2013). For this purpose, an ideal reference gene should be stably and widely expressed in various tissues, samples, or subjects and not be influenced by the experimental condition and treatment. The qPCR primers should also be optimized by tuning a variety of parameters like amplicon length, melting temperature (T_m_), and amplification efficiency (Bustin, 2002; Rebouças et al., 2013).

Researchers commonly choose one or a few reference genes from a set of housekeeping genes stably expressed in the tissues of interest that fit their experimental design. Frequently used reference genes compared and summarized in humans include *beta-actin* (*ACTB1*), *beta-2-microglobulin* (*B2M*), *glyceraldehyde-3-phosphate dehydrogenase* (*GAPDH*), *beta-glucuronidase* (*GUSB*), *hypoxanthine phosphoribosyltransferase 1* (*HPRT1*), *phosphoglycerate kinase* (*PGK1*), *ribosomal protein S19* (*RPS19*), *18S ribosomal RNA* (or *18S rRNA*), *TATA box binding protein* (*TBP*), *beta-tubulin* (*TUBB*), and *ubiquitin C* (*UBC*) (Adeola, 2018; Hellemans and Vandesompele, 2014; Rebouças et al., 2013). Reference genes in mouse samples are often selected from a similar list of housekeeping genes, where *Gaphd*, *Actb1,* and *18S rRNA* are most frequently chosen (Mosley and HogenEsch, 2017; Ruiz-Villalba et al., 2017). Although the RT-qPCR technique was already widely used in the early 2000s, reference gene selection in zebrafish was not well validated until 2007. *18S rRNA*, *actb1*, and *eukaryotic translation elongation factor 1 alpha 1, like 1* (*eef1a1l*) are frequently used in zebrafish (McCurley and Callard, 2008; Tang et al., 2007).

*18s rRNA* genes are highly conserved across all eukaryotes and expressed in all living cells and are, therefore, ideal reference genes across species (Kozera and Rapacz, 2013). Unlike the relatively well-studied mRNA/protein gene-coding regions in human and mouse genomes, the sequences that contain zebrafish *18S rRNA* (*rna18s*) genes were not well-sequenced nor mapped in the genome assembly (Locati et al., 2017), which has made the primer design has been challenging. In this paper, we summarize some incorrect and sub-optimal zebrafish *rna18s* qPCR primers that have been used in quantitative studies of mRNA expression since 2006. In addition, we have designed and tested primer sets to recommend for future studies.

## Results

### 1. A widely but wrongly used primer for zebrafish *rna18s* qPCR

We looked through research papers using *rna18s* as a qPCR reference gene in zebrafish and found that most of them used *rna18s* qPCR primers with errors. The most widespread error in the community is a primer set of 5’-TCGC<u>**ta**</u>GT<u>**t**</u>GGCATCGTTTA<u>**t**</u>G-3’ (referred to as rna18s-F1-W1 in this article, Table 1) and 5’-CGGAGGTTCGAAGACGATCA-3’ (referred to as rna18s-R or rna18s-SS1 in this article, Table 1). Primer rna18s-R matches zebrafish *rna18s* sequence, but rna18S-F1-W1 does not. Four nucleotides (indicated in bold/underlined lowercase letters in rna18s-F1-W1 sequence) are different from the actual zebrafish *rna18s* sequence, 5’-TCGC<u>**GG**</u>GT<u>**C**</u>GGCATCGTTTA<u>**C**</u>G-3’ (mismatched nucleotides indicated in bold/underlined uppercase letters).

**Table 1.**
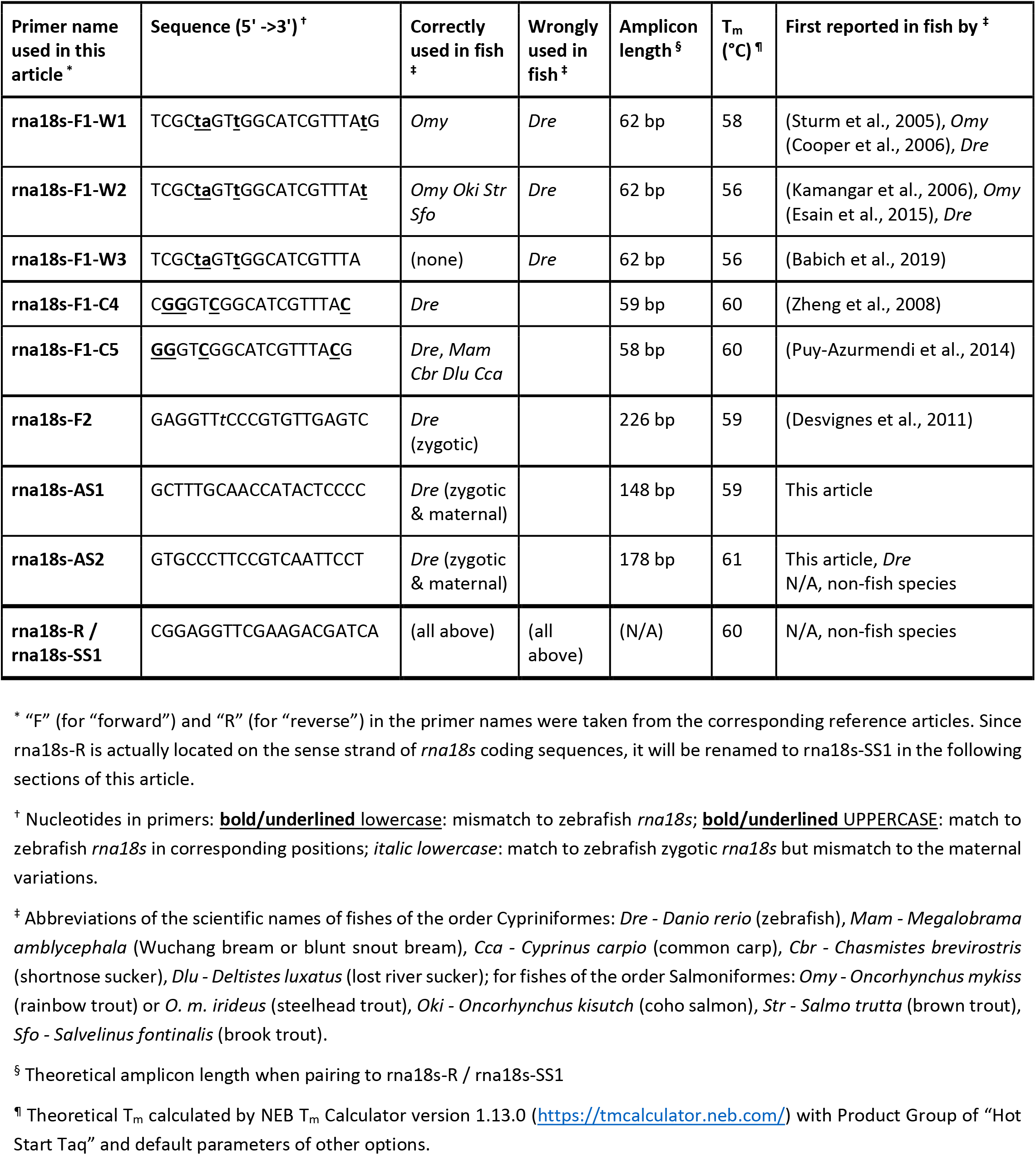
Wrong and correct qPCR primers used to amplify zebrafish rna18s.

Many incorrect usages of rna18s-F1-W1 could be traced back to some key publications specifically analyzing the selection of zebrafish qPCR reference genes (Table 2). One of the most cited articles that used the wrong primers was published by McCurley & Callard in 2008. They incorrectly used rna18s-F1-W1 to amplify zebrafish *rna18s*, the gene they later suggested as a qPCR internal reference in some cases in their conclusion (McCurley and Callard, 2008). As of Aug 2021, this article has been cited more than 500 times. It is hard to find out the precise number of articles that used the wrong *rna18s* RT-qPCR primer set since most of them only cited other publications but did not provide the exact primer sequences they used. However, it is reasonable to assume that a considerable number, if not all, of these publications, followed the McCurley & Callard article and misused rna18s-F1-W1 in zebrafish.

**Table 2.**
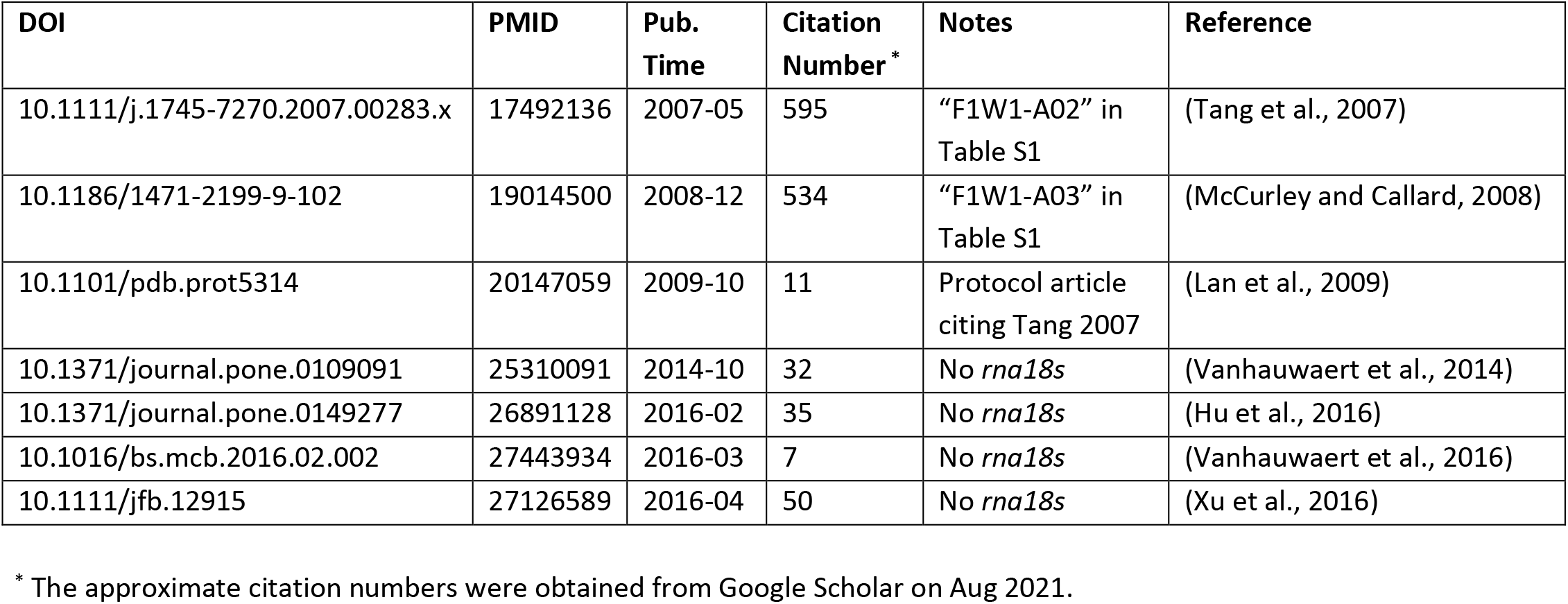
Selected publications comparing zebrafish internal reference genes.

| DOI | PMID | Pub. Time | Citation Number * | Notes | Reference |
| --- | --- | --- | --- | --- | --- |
| 10.1111/j.1745-7270.2007.00283.x | 17492136 | 2007-05 | 595 | "F1W1-A02" in Table S1 | (Tang et al., 2007) |
| 10.1186/1471-2199-9-102 | 19014500 | 2008-12 | 534 | "F1W1-A03" in Table S1 | (McCurley and Callard, 2008) |
| 10.1101/pdb.prot5314 | 20147059 | 2009-10 | 11 | Protocol article citing Tang 2007 | (Lan et al., 2009) |
| 10.1371/journal.pone.0109091 | 25310091 | 2014-10 | 32 | No *rna18s* | (Vanhauwaert et al., 2014) |
| 10.1371/journal.pone.0149277 | 26891128 | 2016-02 | 35 | No *rna18s* | (Hu et al., 2016) |
| 10.1016/bs.mcb.2016.02.002 | 27443934 | 2016-03 | 7 | No *rna18s* | (Vanhauwaert et al., 2016) |
| 10.1111/jfb.12915 | 27126589 | 2016-04 | 50 | No *rna18s* | (Xu et al., 2016) |

\* The approximate citation numbers were obtained from Google Scholar on Aug 2021.

We checked some other articles focusing on comparing zebrafish internal reference genes (Table 2). The Tang article, published slightly earlier to McCurley and Callard’s, also tested several reference genes. They provided primer sequences of all other genes except for *rna18s*, indicating it as “a generic Taqman probe supplied by Applied Biosystems” (Tang et al., 2007). The amplicon length of this primer set is also 62 bp, the same as the rna18s-F1-W1/rna18s-R. Tang’s article also suggested *rna18s* for future qPCR experiments as part of its findings and has been cited by about 600 articles, most of which cited McCurley & Callard’s article as well and might have followed their misusage of rna18s-F1-W1. None of the remaining five articles directly evaluated the *rna18s* gene or provided any *rna18s* primer sequences, whether correct or wrong, and have been cited much fewer times.

In addition, some other articles provided the primer sequences directly without any citation. The lack of full-text academic searching tools and the academic journal paywalls limited our ability to find all articles providing the primer sequences. We used Google Scholar to look for articles explicitly claimed to have used the primer rna18s-F1-W1 by searching its sequence and then manually checking and classifying each result entry. As of Aug 2021, we have found 81 entries from the Google Scholar search results, including 60 academic articles, 19 theses, and 2 duplicated entries (Supplementary Table S1, publications numbered as “F1W1-01” to “F1W1-81”). Among the 60 peer-reviewed (from journals) or pre-print (from www.bioRxiv.org) articles, 50 used this primer set wrongly on zebrafish cDNA samples, 8 used it correctly on rainbow trout samples, and 2 used it correctly on non-fish species. Furthermore, we found more publications that were not included in the Google Scholar searching results after tracing back to the source of qPCR primers in each article (Supplementary Table S1, publications referred to as “F1W1-A01” to “F1W1-A13”). The two most cited key publications previously mentioned focusing on the comparison of zebrafish RT-qPCR reference genes, namely the Tang article and the McCurley & Callard article were included in the extended list (referred to as “F1W1-A02” and “F1W1-A03” in Supplementary Table S1, respectively).

After tracing further back to the origin of this misusage of rna18s-F1-W1 in zebrafish, we found the primer set was initially designed based on the *rna18s* sequence of rainbow trout (*Oncorhynchus mykiss*), a fish species of the order Salmoniformes, instead of zebrafish (*Danio rerio*), a fish of the order Cypriniformes. This primer set was correctly used in rainbow trout samples by a group from Institut Fedératif de Recherche, France (INFR) and King’s College London, UK (KCL) at first (Sturm et al., 2005) (referred to as “F1W1-14” in Supplementary Table S1) and continued to be used in the same species by the rainbow trout community in the following years. In 2006, this primer set was first reported to be used on zebrafish samples in an article by another group from KCL with overlapping co-authors with the previous one on rainbow trout, even though the primer sequences do not fully match the zebrafish sequences (Cooper et al., 2006) (referred to as “F1W1-A01” in Supplementary Table S1). The latter group also claimed that this primer set yields 150 bp amplicons from zebrafish total RNA, while the actual product should be 62 bp in both rainbow trout and zebrafish if the primer anneals to the predicted location. Later, one of the two mentioned key zebrafish reference gene publications, McCurley & Callard article, used this wrong *rna18s* primer set, citing the second KCL group, and then recommended it to the zebrafish community in 2008 (McCurley and Callard, 2008). The sequences or citations of the rna18s primer set of the other key publication, the Tang article, were not provided (Tang et al., 2007).

Moreover, we found two more primers based on rainbow trout or other Salmoniformes fish genomes or RNA sequences but used on zebrafish samples. These two primers are highly similar to rna18s-F1-W1 with only one or two nucleotides removed from the 3’-end (referred to as rna18s-F1-W2 and rna18s-F1-W3 in this article, representativity; Table 1 and Figure 1). Primer rna18s-F1-W2 lacks the last nucleotide of rna18s-F1-W1, leaving the first nucleotide at the 3’-end mismatched to the zebrafish template, which might theoretically affect the PCR amplification when using these samples as the templates. This primer was found in 27 articles via Google Scholar searching (Supplementary Table 1, publications referred to as “F1W2-01” to “F1W2-27). Of the 27 articles, 26 correctly used the primer on Salmoniformes fish samples, while the remaining one incorrectly used it on zebrafish samples (Esain et al., 2015). This primer was found to be used in two earlier zebrafish articles by the same group published in 2016 (Kwan et al., 2016) and 2020 (Lundin et al., 2020), which were not included in the Google Scholar search results (Supplementary Table 1, publications referred to as “F1W2-A01” and “F1W2-A02”).

**Fig. 1.**
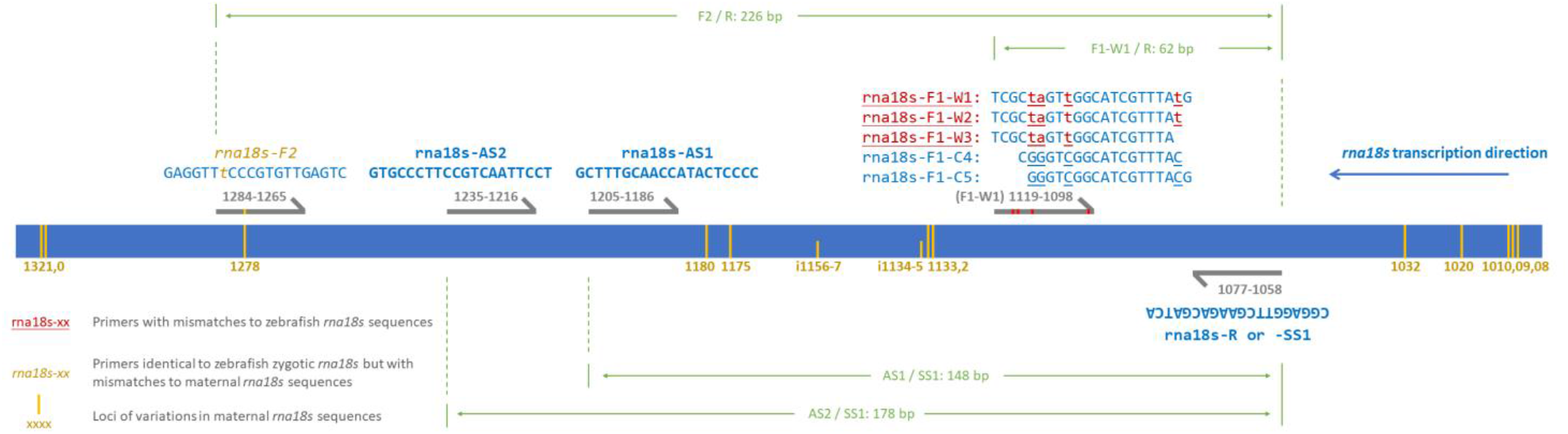
Sequences of qPCR primers reviewed and newly designed in this article and their locations on zebrafish zygotic *rna18s* RNA transcript.

Primer rna18s-F1-W3 lacks the last two nucleotides of rna18s-F1-W1. It contains 11 continuous identical nucleotides at the 3’-end and all three mismatches at the 5’-parts when used in qPCR amplifying zebrafish templates, which might theoretically slightly reduce the mismatch effects (Table 1 and Fig. 1). This primer was only found in 1 article published in 2019 and wrongly used on zebrafish samples (Babich et al., 2019) (Supplementary Table 1, publications referred to as “F1W3-01”).

### 2. Other zebrafish *rna18s* primers with correct sequences at the same region

Other than the wrong primers rna18s-F1-W1, -W2, and W3, we did find two more primers that are based on the correct zebrafish *rna18s* sequence in the same region (will be referred to as rna18s-F1-C4 and rna18s-F1-C5, Table 1 and Fig. 1). Both primers are four nucleotides shorter than rna18s-F1-W1, possibly designed to maintain a proper T_m_.

Primer rna18s-F1-C4 was found in three zebrafish qPCR articles published in 2008-2013 (Supplementary Table S1, publications referred to as “F1C4-01” to “F1C4-03”) by the zebrafish KCL groups mentioned before and other relevant groups from KCL, which was first described as “designed using Primer Express (Applied Biosystems)” (Zheng et al., 2008). Regrettably, this correct zebrafish qPCR primer set, with the accurate description of amplicon length of 62 bp and providing the Taqman probe sequence used together, has not been cited or used by any other groups (at least according to Google Scholar search results), even if it came from the same group that first distributed the rainbow trout sequences incorrectly as a zebrafish primer set.

The sequence of primer rna18s-F1-C5 was shared by zebrafish and some other Cypriniformes fishes. It has been correctly used on samples of different fish species from at least 23 publications (Supplementary Table S1, publications referred to as “F1C5-01” to “F1C5-23”), including 22 journal articles, most of which (16) are on *Megalobrama amblycephala* (Wuchang bream or blunt snout bream), 3 on zebrafish, and 3 on other Salmonidae fishes. The three peer-reviewed zebrafish articles were published by two different groups, one in 2014 (Puy-Azurmendi et al., 2014) and the others in 2015 and 2017 (He et al., 2017; Huang et al., 2015).

### 3. Another zebrafish *rna18s* primer used in zygotic zebrafish sample

We also found a few articles that used another qPCR primer designed to amplify a 226-bp region in zebrafish samples (referred to as rna18s-F2; Table 1 and Fig. 1) when paired with rna18s-R. Of the 11 publications from Google Scholar search results (Supplementary Table S1, publications referred to as “F2-01” to “F2-11”), primer rna18s-F2 was reported by 7 articles, including 4 used on zebrafish research, one of which was from a group of INRA (Desvignes et al., 2011) and the other three were from another group of in the same institute. The amplicon length of the rna18s-F2/rna18s-R primer set (226 bp) is longer than that of the rna18s-F1’s/rna18s-R pairs (58 to 62 bp, Table 1 and Fig. 1). It is also longer than the typical suggested length when using Taqman systems (50-150 bp) or SYBR systems (70-200 bp), which might influence qPCR efficiency (Debode et al., 2017).

A critical point to note when using the rna18s-F2/rna18s-R primer set is that *rna18s* RNA transcripts are encoded at multiple genomic loci in zebrafish. Some of these loci have been mapped and annotated in the current zebrafish genome assembly (GRCz11), but others have not. Some of the mapped *rna18s*-encoding sequences were studied for their nucleotide variations and expression profiles in different development stages. The predominate maternal or zygotic variants in zebrafish are not identical (Locati et al., 2017). The primers of rna18s-R and rna18s-F1 (correct versions) anneal to a conserved region of the target *rna18s* sequence. However, the sequence of rna18s-F2 is identical to the zygotic zebrafish *rna18s* sequences but contains a mismatch to the maternal RNA variants, which are the main components of zebrafish RNA samples before the 128-cell stage or 4.3 hours post-fertilization (Fig. 1). All four zebrafish articles using rna18s-F2/rna18s-R have applied this primer set only in zygotic samples. No analysis of qPCR results by this primer pair in early-staged zebrafish samples was reported, and we do not know whether this mismatch would affect the amplification in all cases.

### 4. Other incorrect zebrafish *rna18s* primer usage or information provided

We also found other types of errors in articles claiming to use zebrafish *rna18s* as qPCR reference genes. The first error is amplicon size discrepancy. As we mentioned, the KCL group provided an amplicon length of 150 bp in their first article reported the usage of rna18s-F1-W1/rna18s-R in zebrafish samples (Cooper et al., 2006), despite that the correct length should be 62 bp if the amplification worked as expected when employing Taqman qPCR systems. The amplicon size with the same primer set was reported as a variety of sizes from 85 bp to 123 bp in other articles. All these cases were collected in columns “H” of Supplementary Table S1.

We found another common error: some articles confused *rna18s* (ZFIN ID: ZDB-RRNAG-180607-2) with other genes. Palstra et al. provided the primer sequences of rna18s-F1-W1/rna18s-R but wrongly labeled them as *rps18*, the symbol of *ribosomal protein S18* (ZFIN ID: ZDB-GENE-020419-20), a protein-coding gene whose products bind to rRNAs and participate in the formation of the ribosome complexes (Palstra et al., 2010). In another article, Tzung et al. provided the rna18s-F1-W1/rna18s-R sequences with the label *“18s”* in their original reports (Tzung et al., 2015a). Therefore, in a correction note, the authors revised the primer sequences to another pair identical to *rps18* sequences but kept the gene symbol label unchanged (Tzung et al., 2015b). Some articles did not provide the primer sequences with the gene symbols but chose to cite a GenBank or RefSeq ID instead. However, not all database IDs correctly represented zebrafish *rna18s*. For example, Yu et al. claimed their *rna18s* primers were designed based on RefSeq sequence NM_100073, which was the sequence ID of zebrafish gene *rpl18* (*ribosomal protein L18*, ZFIN ID: ZDB-GENE-040801-165), another protein-coding gene that encodes products related to ribosome complexes; and the actual materials they used were unclear (Yu et al., 2010). These cases of incorrect gene names, gene symbols, or database IDs were collected in columns “G” of Supplementary Table S1.

Another issue we have seen is the questionable usage of materials in the reverse transcription (RT) steps. As an RNA gene that does not code for any protein, *rna18s* is a part of the transcript complex of *45S rRNA* transcribed by RNA polymerase I. The rRNA transcripts lack poly(A) tails, different from the mRNA transcripts of most protein-coding genes transcribed by RNA polymerase II and then processed by capping and tailing steps (Gerbi, 1995). Non-poly(A)-tailed transcripts could only be reversed transcribed by adding randomized nucleotide primers, such as random hexamers provided by the manufacturer of the kits or elsewhere. Despite these facts, several articles claimed that they only used oligo d(T)s as the RT primers to synthesize the first cDNA strands, including *rna18s* from total RNA samples, which is theoretically inappropriate. The information of RT primers, if provided, was collected in columns “I” of Supplementary Table S1.

### 5. New RT-qPCR primer design and efficiency comparison

When paring with the rna18s-R, rna18s-F1-C4 and rna18s-F1-C5 correspond to amplicon lengths of 59- and 58-bp while rna18s-F2 correspond to 226-bp, neither falls in the range of typical amplicon length when using SYBR systems (70-200 bp) (Debode et al., 2017). As was mentioned above, rna18s-F2 has mismatches when used on zebrafish maternal RNA samples. To find alternative qPCR primer sets without these potential defects, we first used a pair of outer primers to amplify a region containing all reported qPCR primers. Sanger sequencing results showed that the actual sequences of *rna18s* (zygotic) of zebrafish were identical to sequences between 165,514 and 165,758 of GenBank entry BX296557.35 (Supplementary Fig. S1), which confirmed the template sequences. Two new qPCR primers at the anti-sense strand of non-variation regions of maternal & zygotic *rna18s* transcripts, rna18s-AS1 and rna18s-AS2, were designed by pairing with the common primer rna18s-R on the sense strand (renamed to rna18s-SS1 here). The amplicon lengths of the new primer sets were 148 bp and 178 bp, respectively (Table 1 and Fig. 1).

We parallelly tested five primers sets. All sets contained the same sense primer (rna18s-SS1, i.e., rna18s-R) and different anti-sense primers, including three previously described ones: rna18s-F1-W1, rna18s-F1-C4, and rna18s-F2, as well as two newly designed ones: rna18s-AS1 and rna18s-AS2. All primers had a similar T_m_ (Table 1). The same template samples in 6-point 10x serial dilutions (1 to 1×10^−5^ x) were used in SYBR qPCR systems with the same programs. Our results showed that all four primer sets without mismatches could get standard curves with R^2^ > 0.995, amplification efficiencies between 91% to 99%, and C_q_ values of all six concentration data points ranging from 11-33. However, for the mismatched primer set (rna18s-F1-W1/rna18s-R), the C_q_ values of the four lowest cDNA concentration points were too high to be regarded as valid. Only C_q_ values about 35-40 of the remaining two high concentration points were obtained, and the standard curve could not be considered valid (Fig. 2). The results suggested the mismatched primer set is low in efficiency and specificity, and the two new optimized primers were suitable for qPCR experiments.

**Fig. 2.**
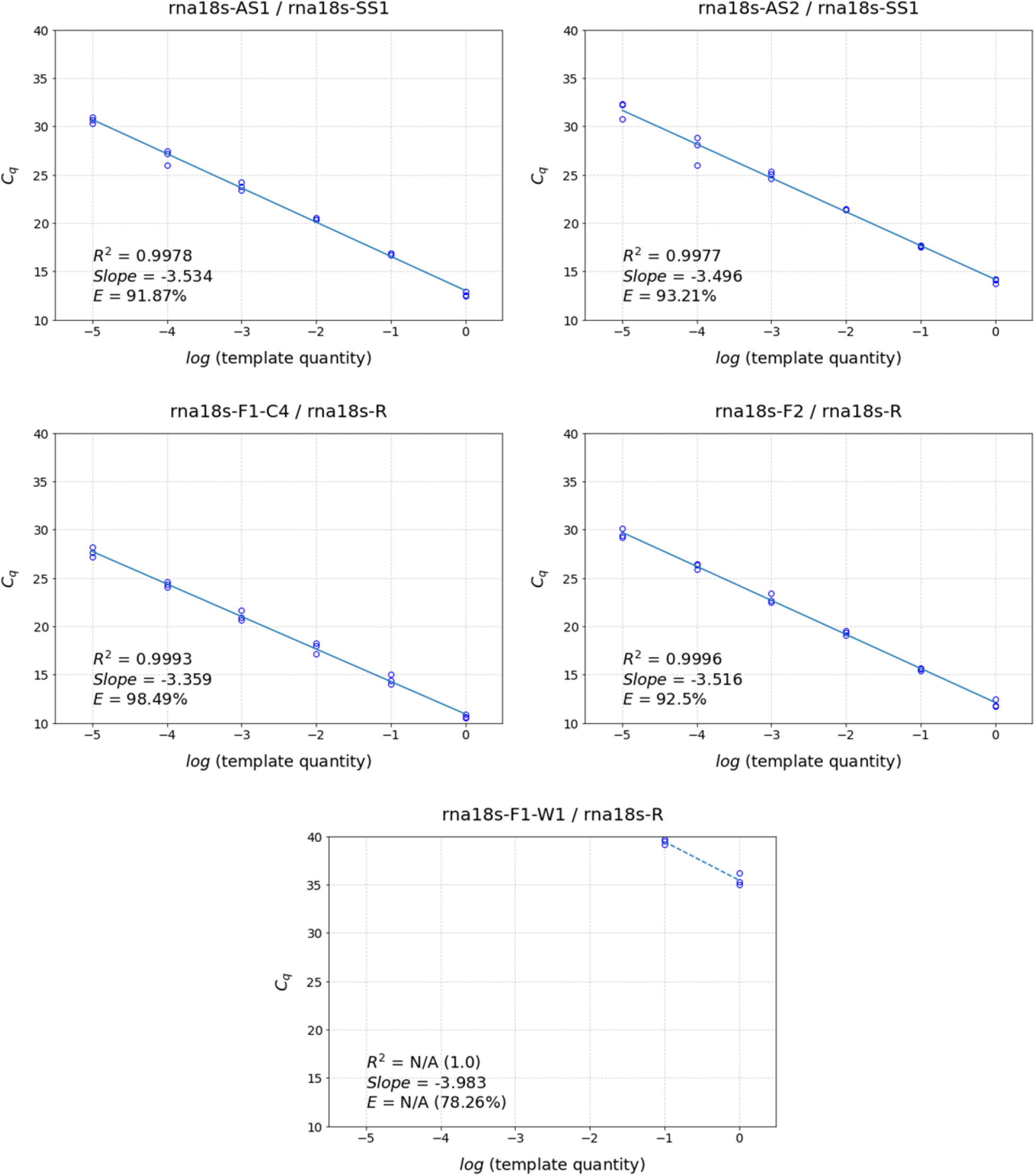
C_q_ values and standard curves of tested primer sets. The standard curve of the primer set rna18s-F1-W1 / rna18s-R can not be considered valid with only two data points. For this primer set, assumed R2 and E are calculated for comparison.

## Discussion

*18S rRNA* was wildly used as a qPCR reference gene in many species. It may be due to its high conservation across species and high expression level across cells and tissues. Ribosomal RNAs (rRNAs) are the most abundant form in the total RNA of cells. *18S rRNA* is the main component of the eukaryotic ribosomal small subunit, essential for translation initiation and mRNA-tRNA interaction. It is universally demanded in spatiotemporal cycles of all living cells and is believed to be a relatively stably expressed gene. However, like other RNA genes, the *18S rRNA* gene is different from the protein-coding mRNA genes in many ways. At the transcriptional level, *28S*, *5.8S*, and *18S rRNAs* are processed from a *45S rRNA* transcription unit, lacking 5’-cap or 3’-poly(A) tail structures. At the genomic level, hundreds of *45S rRNA* encoding unit copies are organized in tandem repeats and present in clusters located in different genome regions. These characteristics may cause them to have additional considerations when used as reference genes. For example, the reverse-transcription primers to synthesize the first strand of cDNA should be carefully selected.

One of the challenges of designing primer sets for rRNA genes is the difficulty of finding accurate and detailed information in bioinformatic databases compared to mRNA/protein-coding genes. In many species (e.g., zebrafish), few of the sequences and genomic loci of rRNA genes are well determined. The same rRNA transcript is often encoded at multiple chromosomal loci, which could be poorly mapped on the assembled genome, containing a lot of variations even in the same individual. The three authorized gene nomenclature consortiums-concerned hub gene databases, HGNC, MGI, and ZFIN, contain entries of protein-coding genes of the three model animals, human, mouse, and zebrafish, respectively. They are also the founding members of Genome Reference Consortium (GRC), which maintains the current genome assemblies (Church et al., 2011). However, they document rRNA genes in various styles with significant differences to protein-coding genes. It is known that there are five main rRNA-encoding clusters located at five chromosome regions in the human genome, but the exact assembly positions have not yet been finalized in the current human genome assembly GRCh38. Each cluster contains hundreds of times tandem repeated *45S rRNA* units. The known human *45S rRNA* clusters (*RNR1* to *RNR5*; HGNC IDS: 10082-10084) and their *18S rRNA* subgroups (with grouped symbols *RNA18S1* to *RNA18S5*; HGNC IDs: 44278-44281, and 37657) are well organized by HGNC (see https://www.genenames.org/data/genegroup/#!/group/1381). On the other hand, mouse rRNA genes were not fully annotated or named. Currently, there are eight annotated mouse *Rna18s* genes or gene clusters in MGI: *Rn18s*, *Rn18s-rs1* to *Rn18s-rs7* (MGI IDs: 97943 and 107310, 107305, 107276, 107275, 107222, 107219, 107213), of which *Rn18s* and *Rn18s-rs5* have been mapped to specific loci but others are only associated with chromosomes.

In the case of zebrafish, rRNA genes have been much worse studied. ZFIN collects all zebrafish *rna18s* genes under one unique entry (ZFIN ID: ZDB-RRNAG-180607-2; see http://zfin.org/ZDB-RRNAG-180607-2), with no chromosome mapping information, leaving the total number of rRNA clusters unknown. Many articles cite GenBank entry ID BX296557 (Zebrafish DNA sequence from clone CH211-284O1 in linkage group 20, complete sequence, see https://www.ncbi.nlm.nih.gov/nuccore/BX296557) as the reference sequence of zebrafish *rna18s* gene, which is a 173,360-bp large BAC clone containing at least one cluster of *45S rRNA* units. This sequence entry has been updated up to 35 times in two years from 2003 to 2005, and the tandem repeat numbers and exact sequences of rRNA units changed frequently between the versions. The BAC clone itself is also not stably placed in the same place in Chromosome 20 in different zebrafish genome assemblies. Bioinformatics database like RNAcentral has tried to collect as much non-coding RNA information as possible (RNAcentral Consortium, 2021). The sequence variations between zygotic and maternal *rna18s* transcripts were well analyzed and described in detail at the transcript level as components of the zebrafish *45S rRNA* transcripts (Locati et al., 2017). However, reliable annotated sequences of rRNA genes in the zebrafish genome are still in need, contributing to the chaos and misusage of zebrafish *rna18s* qPCR primers.

According to our retrospective work, zebrafish *rna18s* was mainly suggested as a qPCR reference gene by two top-cited articles to the community. The Tang article evaluated several candidate qPCR reference genes in zebrafish. They found that *rna18s* (*18S rRNA*) only showed moderate expression stability in different developmental stages compared to *actb1* (named initially as *β-actin*), *rpl13a*, and *eef1a1l1* (*ef1α*), but was very stable in different tissue types similar to *rpl13a* and *eef1a1l1* (Tang et al., 2007). The McCurley & Callard article compared the stability of expression profiles of several zebrafish housekeeping genes, including *rna18s*. They found *rna18s*, *b2m*, and *eef1a1l1* (*elfa*) had the most stable expression across sexes, developmental stages, and tissue types. In contrast, *eef1a1l1*, *actb1* (*bactin1*), and *tuba1b* (*tuba1*) expression were resistant to the effects of hormone or toxicant treatments (McCurley and Callard, 2008). As part of their conclusion, *rna18s* is recommended as a zebrafish qPCR reference gene in some conditions. However, the *rna18s* qPCR primer set they used was either wrong (i.e., rna18s-F1-W1) or unclear, and thus the conclusions concerning *rna18s* need to be used with caution as references.

In summary, further use of the wildly used primer rna18s-F1-W1, which was designed for rainbow trout, should be avoided in zebrafish. To perform perfect amplification of zebrafish *rna18s*, choosing correct qPCR primer sets is essential. We strongly recommend that the following points be considered if researchers want to design their own *rna18s* primer sets: the amplicon lengths for relevant qPCR systems (Taqman, SYBR, etc.) the variation between zygotic and maternal sequences. Four tested primer sets are recommended by this article with the same sense-strand primer rna18s-SS1 (rna18s-R), pairing with different anti-sense-strand primers to get amplicons with different lengths: rna18s-AS2 (178 bp), rna18s-AS1 (148 bp), rna18s-F1-C4 (59 bp), and rna18s-F1-C5 (58 bp). They could be considered in zebrafish *rna18s* qPCR assays with diverse needs.

## Methods and Materials

### Summarization of Google Scholar search results and the extended publication lists

Raw sequences of the rna18s-F primers were directly searched in Google Scholar (https://scholar.google.com/) in Aug 2021. Each result entry was checked, collected, and sequentially numbered as “<primer name>-xx”. The information of publication DOI’s or URL links (if DOI not available), as well as the PMID and the first publish time, were recorded if they were available. Publications were categorized by types (article, book, thesis, or other) and the species of samples on which the primer set was used. For zebrafish articles, the claimed citation of publications and database IDs of *rna18s* primer sequences were recorded, as well as the reverse transcription primers and Taq probe sequences they claimed to use, if provided. Lengths of qPCR amplicons were only recorded if they were different from the theoretical values. Publications not listed in the Google Scholar search results but cited by any articles listed were also collected in a similar style and sequentially numbered as “<primer name>-Axx”.

### Sample harvesting and total RNA extraction

TUxAB wild-type zebrafish larvae were collected at the stage of 7 days post fertilization (dpf). 20 larvae were put together into tubes containing 500 μL TRIzol (Invitrogen 10296010) and frozen at −80°C before the next steps. RNA extraction was done following the manufacturer’s protocol. Briefly, each frozen sample tube was thawed and warmed up to room temperature (RT) before 500 μL TRIzol was added to get a final volume of 1 mL. 200 μL of chloroform was then added to each sample, gently mixed, and incubated at room temperature (RT) for 2-3 minutes, followed by centrifugation at 4°C, 12,000x g for 15 minutes. The upper aqueous phase was transferred to a new tube, and 500 μL isopropanol was added and mixed well for RNA precipitation. After centrifuging at 4°C, 12,000x g for 30 minutes, the supernatant was discarded, and the RNA pellet was washed with 70% ethanol once, air-dried, and then resuspended in 20 μL of RNase-free water. 1 μL rDNase I and 2 μL DNase I Buffer from DNase Treatment and Removal Reagents Kit (ThermoFisher Scientific AM1906) were added to each RNA sample and then incubated at 37°C for 30 min. 2 μL of DNase Inactivation Reagent was added and incubated at 37°C for 2 min. The RNA samples were then quantified using NanoDrop spectrophotometers.

### Reverse transcription and template preparing

The first strand cDNAs were reverse transcribed using SuperScript IV VILO Master Mix kit (Invitrogen, 11756500) following the manufacturer’s protocol. Briefly, 1 μg of total RNA was used in a 20-μL reaction system, along with both oligo (dT)18 and random hexamer primers in the master mix from the kit. The reaction mixture was incubated at 25°C for 10 minutes, 50°C for 10 minutes, 55°C for 5 minutes, and then mixed with 180 μL water to a final volume of 200 μL to be used as the 1x PCR template. A 10-fold serial dilution was done to prepare 0.1x to 10^−5^x samples, resulting in a total of six concentrations of the template.

### Confirmation of template sequences

The reverse transcription reaction mixture was used for PCR by an outer primer pair with the sequences 5’-TAAGCCGCAGGCTCCACTCC-3’ (identical to GenBank entry BX296557.35, sequence region 165,514 to 165,533) and 5’-GCGCCGCTAGAGGTGAAATTCT-3’ (identical to BX296557.35, sequence region 165,787 to 165,808). A unique amplicon of around 300 bp was obtained. The PCR product was purified and sent for Sanger sequencing with both outer primers. Sequencing results were aligned to the sequence of GenBank entry BX296557.35 for analysis.

### Designing of new qPCR primers

New optimized primers for SYBR qPCR system were designed by NCBI Primer-BLAST (with Primer3 program in the background) by providing the template sequences as the region of zebrafish zygotic *rna18s* confirmed in the previous step. Default parameters were used except for the following items: “Use my own forward primer (5′->3′ on plus strand)”: CGGAGGTTCGAAGACGATCA (i.e. the sequence of rna18s-R or rna18s-SS1); PCR product size (bp): 70-200; primer melting temperatures (T_m_, °C): min 55, opt 60 max 65, max difference 3; primer size (bp): min 18, opt 20, max 30; primer GC content (%): min 40.0, max 60.0; repeat filter: none. Two anti-sense primers were selected from the result lists without variations between zebrafish zygotic and maternal *rna18s* sequences.

### RT-qPCR of reported and newly designed primer sets

Parallel qPCR reactions of the six template concentrations were run in triplicate for each primer set in a 384-well plate. In each well, a 10-μL reaction system was prepared, containing 2 μL template sample, 3 μL of 1 μM primer mix, and 5 μL of 2x iTaq Universal SYBR Green Supermix (Bio-rad, 1725124). A two-step qPCR program was employed: 94°C 2 min, (94°C 20 s, 60°C 30 s) x 45 cycles, followed by melting curve analysis: 60 to 95°C (increments of 0.5°C/2 s).

