## Supplementary Fig. S1 for "Double-check the zebrafish *18s rRNA* qPCR primers: they may be wrong"

### Reference sequence of zebrafish zygotic *rna18s*

**(GenBank: BX296557.35, about 165,900 to 165,500)**

*rna18s* transcription direction
(reverse complementary of BX296557.35) →

outer primer #1 for PCR & sequencing ->

.. GGGTTTC*t*GGAACCCGGGGCCATGAT*t*GAGAGGGACGGCCGGGGGCATTCGTATTGCGCCGCTAGAGGTGAAATTCTTGGACCGGCGCAAGACGGAC*gaa*AGCGAAAGC*a*TTTGCCAAGA

rna18s-R / rna18s-SS1 ->

A*t*GTTTTCATTAATCAAGAACGAAAGTCGGAGGTTCGAAGACGATCAGATACCGTCGTAGTTCCGACCGTAAACGATGCCGACCCGCGATCCGGCGGCGTT*at*T^CCCATGACCCGCCGGG

<- rna18s-F1 (5 versions)

CAGCGT^GCGGGAAACCACGAGTCT*t*TGGG*t*TCCGGGGGGAGTATGGTTGCAAAGCTGAAACTTAAAGGAATTGACGGAAGGGCACCACCAGGAGTGGAGCCTGCGGCTTAATTTGACTCA

<- rna18s-AS1 <- rna18s-AS2 <- outer primer #2 for PCR

ACACGGG*a*AACCTCACCCGGCCCGGACACGGA ..

<- rna18s-F2

*n* ^ loci of variations in maternal *rna18s* sequences, as described by Locati et al. 2017

NNN zygotic *rna18s* sequence region confirmed by PCR & Sanger sequencing results

### Sanger sequencing results example

**(sequenced by outer primer #1)**

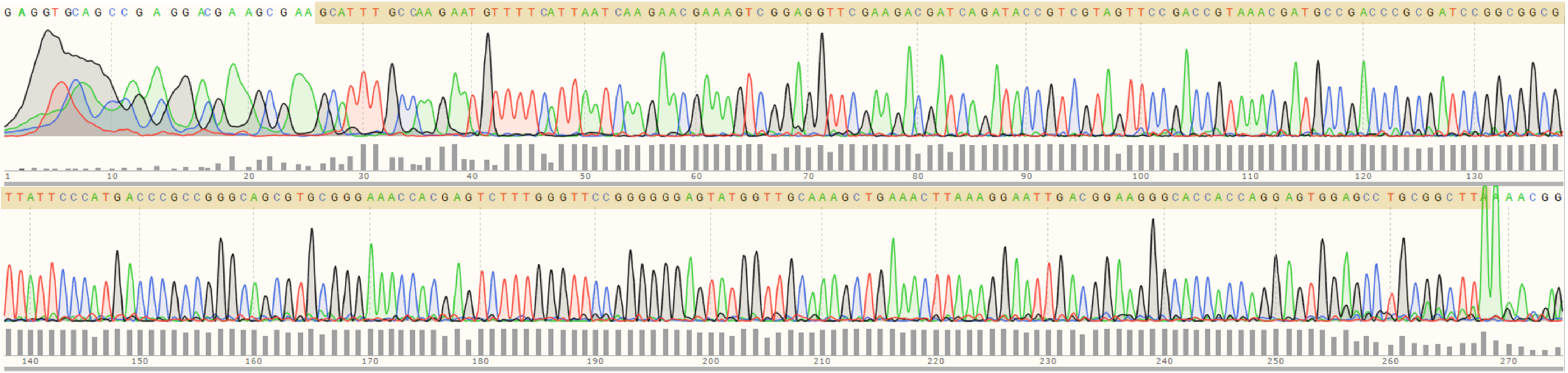
